# Tomato wall-associated kinase SlWak1 acts in an Fls2- and Fls3-dependent manner to promote apoplastic immune responses to *Pseudomonas syringae*

**DOI:** 10.1101/2020.01.27.921460

**Authors:** Ning Zhang, Marina A Pombo, Hernan G Rosli, Gregory B. Martin

## Abstract

Wall-associated kinases (Waks) are known to be important components of plant immunity against various pathogens including *Pseudomonas syringae* pv. tomato (*Pst*) although their molecular mechanisms are largely unknown. In tomato, *SlWak1* has been implicated in immunity because its transcript abundance increases significantly in leaves after treatment with the flagellin-derived peptides flg22 and flgII-28, which activate the receptors Fls2 and Fls3, respectively. We generated two *SlWak1* tomato mutants (Δwak1) using CRISPR/Cas9 and investigated the role of *SlWak1* in tomato-*Pst* interactions. PTI activated in the apoplast by flg22 or flgII-28 was compromised in Δwak1 plants but PTI at the leaf surface was unaffected. The Δwak1 plants developed fewer callose deposits than wild-type plants but retained the ability to generate reactive oxygen species and activate MAPKs in response to flg22 and flgII-28. The induction of *Wak1* gene expression by flg22 and flgII-28 was greatly reduced in a tomato mutant lacking Fls2 and Fls3 but induction of *Fls3* gene expression by flgII-28 was unaffected in Δwak1 plants. After *Pst* inoculation, Δwak1 plants developed disease symptoms more slowly than Δfls2.1/fls2.2/fls3 mutant plants, although both plants ultimately were similarly susceptible. SlWak1 co-immunoprecipitated with both Fls2 and Fls3 independently of flg22/flgII-28 or Bak1. These observations suggest that SlWak1 acts in a complex with Fls2/Fls3 and plays an important role at later stages of the PTI in the apoplast.

## Introduction

Plants have evolved a sophisticated, two-layered inducible defense system, consisting of pattern-recognition receptor (PRR)-triggered immunity (PTI) and NOD-like receptor (NLR)-triggered immunity (NTI), to protect themselves against infection by pathogenic microbes (Jones & Dangl, 2006; Zipfel, 2014). To initiate the PTI response, host PRRs detect potential microbial pathogens by recognizing diverse microbe/pathogen-associated molecular patterns (MAMPs or PAMPs) including peptides from bacterial flagellin (Felix *et al*., 1999). The resulting PTI responses include the production of reactive oxygen species (ROS), activation of mitogen-activated protein kinase (MAPK) cascades, callose deposition at the cell wall, transcriptional reprogramming of immunity-associated genes, and moderate inhibition of pathogen growth (Chandra *et al*., 1996; Jia & Martin, 1999; Zipfel, 2014; Li *et al*., 2016). Two PRRs, Fls2 and Fls3, bind the flagellin-derived MAMPs flg22 and flgII-28, respectively, and in concert with the co-receptor Bak1 (in tomato, Serk3A and/or Serk3B) activate intracellular immune signaling (Chinchilla *et al*., 2007; Sun *et al*., 2013; Hind *et al*., 2016).

To overcome PTI, pathogens deliver virulence proteins (effectors) into the plant cells to interfere with MAMP detection or PTI signaling and promote disease development (Dou & Zhou, 2012). AvrPto and AvrPtoB, two effectors from *Pseudomonas syringae* pv. tomato (*Pst*), suppress the early PTI response by interfering with the interaction of Fls2 with Bak1 (Xiang *et al*., 2008; Martin, 2012; Hind *et al*., 2016). In response to bacterial effectors, plants have evolved genes encoding NLRs (nucleotide-binding oligomerization domain-like receptors) which recognize specific effectors and activate NTI (Martin *et al*., 2003; Jones & Dangl, 2006). In tomato, the Pto kinase protein interacts with AvrPto or AvrPtoB and forms a complex with the NLR protein Prf resulting in the induction of NTI and inhibition of pathogen growth (Martin *et al*., 1993; Salmeron *et al*., 1996; Pedley & Martin, 2003).

Plant cell wall-associated kinases (Wak) or Wak-like kinases (Wakl) are receptor-like protein kinases consisting of an extracellular domain with conserved epidermal growth factor (EGF) repeats, a transmembrane domain, and a cytoplasmic serine/threonine protein kinase domain (Anderson *et al*., 2001). While some Wak proteins play a vital role in cell expansion and plant development (Lally *et al*., 2001; Wagner & Kohorn, 2001; Kohorn *et al*., 2006), others are expressed only in specific organs and differentially regulated by a variety of biotic or abiotic stimuli including pathogen attack (Hou *et al*., 2005; Li *et al*., 2009; Brutus *et al*., 2010; Hu *et al*., 2014; Zuo *et al*., 2015; Lou *et al*., 2019). Wak proteins have been reported to be involved in host resistance against various pathogens in plants including Arabidopsis (Brutus *et al*., 2010), *Nicotiana benthamiana* (Rosli *et al*., 2013), rice (Li *et al*., 2009; Hu *et al*., 2014; Delteil *et al*., 2016; Harkenrider *et al*., 2016), maize (Hurni *et al*., 2015; Zuo *et al*., 2015; Yang *et al*., 2019), and wheat (Yang *et al*., 2014; Saintenac *et al*., 2018; Dmochowska-Boguta *et al*., 2020). In one case, the wheat *Snn1*-encoded Wak protein acts as a susceptibility factor to promote infection of a fungal pathogen *Parastagonospora nodorum* (Shi *et al*., 2016).

Although Wak proteins have been identified as important contributors to disease resistance against various pathogens (Hu *et al*., 2017; Bacete *et al*., 2018), much remains to be learned about the molecular mechanisms they use to activate immune responses. The best-studied Wak protein, the Arabidopsis AtWAK1, recognizes cell wall-derived oligogalacturonides (OGs) and activates OG-mediated defense responses against both fungal and bacterial pathogens (Brutus *et al*., 2010; Gramegna *et al*., 2016). In maize, the ZmWAK-RLK1 protein (encoded by *Htn1*) confers quantitative resistance to northern corn leaf blight (NCLB) by inhibiting the biosynthesis of secondary metabolites, benzoxazinoids (BXs), that suppress pathogen penetration into host tissues (Yang *et al*., 2019). Another ZmWAK protein located in a major head smut quantitative resistance locus *qHSR1* enhances maize resistance to *Sporisorium reilianum* by arresting the fungal pathogen in the mesocotyl (Zuo *et al*., 2015). One wheat Wak protein encoded by the *Stb6* gene recognizes an apoplastic effector (AvrStb6) from *Zymoseptoria tritici* and confers resistance to the fungal pathogen without a hypersensitive response (Saintenac *et al*., 2018). In rice, three OsWAKs act as positive regulators in resistance to the rice blast fungus by eliciting ROS production, activating defense gene expression, and recognizing chitin by being partially associated with the chitin receptor CEBiP (Delteil *et al*., 2016). Wak proteins therefore appear to exhibit extensive functional diversity and have different mechanisms to defend against pathogen infection in different plant species. The functional characterization of Wak proteins in tomato has not been reported and their possible contributions to PTI or NTI are not well understood in this species.

Tomato is an economically important vegetable crop throughout the world and its production is threatened by many pathogens including *Pseudomonas syringae* pv. tomato which causes bacterial speck disease and can result in severe crop losses (Jones, 1991; Kimura & Sinha, 2008). Understanding the functions of Wak proteins in tomato could therefore provide fundamental information for breeding tomato cultivars that are resistant to various pathogens. Tomato contains seven *Wak* and sixteen *Wakl* genes (Zheng *et al*., 2016). The *SlWak1* (Solyc09g014720) gene is clustered together with another three *SlWak* genes (Solyc09g014710, Solyc09g014730 and Solyc09g014740) on chromosome 9; however, the expression of only the *SlWak1* gene (hereafter *Wak1*) is significantly induced after MAMP treatment or *Pst* inoculation (Rosli *et al*., 2013). Knock down of *Wak*1 gene expression in *N. benthamiana* leaves using virus-induced gene silencing (VIGS) compromised resistance to the bacterial pathogen *Pst*. However, three closely-related *NbWak* genes were simultaneously silenced in these experiments, making it unclear if one or a combination of *NbWak* genes contributed to the enhanced susceptibility to *Pst* (Rosli *et al*., 2013). To gain a deeper insight into the role of *Wak1* in tomato-*Pst* interactions, we generated two homozygous *Wak1* mutant lines (Δwak1) in tomato using CRISPR/Cas9. Characterization of these Δwak1 mutants indicated that Wak1 protein acts as an important positive regulator in later stages of flagellin-mediated PTI response in the apoplast and associates in a complex with Fls2 and Fls3 to trigger immune signaling.

## Methods and Materials

### Generation of *Wak1* tomato mutants using CRISPR/Cas9

To mutate the *Wak1* gene in tomato, we designed two guide RNAs (Wak1-gRNA1: GTTAAGATTAGCATAAAACA; Wak1-gRNA2: GGGGCGGTGGCATTCGTTGG) targeting the first exon of *Wak1* using the software Geneious R11 (Kearse *et al*., 2012). Each gRNA cassette was cloned into a Cas9-expressing binary vector (p201N:Cas9) by Gibson assembly as described previously (Jacobs *et al*., 2017). Tomato transformation was performed at the biotechnology facility at the Boyce Thompson Institute. *Agrobacterium* cells containing each gRNA/Cas9 construct were pooled together and used for transformation into the tomato cultivar Rio Grande (RG)-PtoR, which has the *Pto* and *Prf* genes. To determine the mutation type, genomic DNA was extracted from cotyledons or young leaves of each transgenic plant using a modified CTAB method (Murray & Thompson, 1980). Genomic regions spanning the target site of the *Wak1* gene were amplified with specific primers (**Supplemental Table S1**) and sequenced at the Biotechnology Resource Center (BRC) at Cornell University. Geneious R11 and the web-based tool called Tracking of Indels by Decomposition (TIDE; https://tide.deskgen.com) (Brinkman *et al*., 2014) were used to determine the mutation type and frequency using the sequencing files (ab1. format) as described (Zhang *et al*., 2020).

### Off-target evaluation

To investigate potential off-target mutations caused by gRNAs in the Δwak1 plants, Wak1-gRNA1, which induced target mutations in *Wak1* in the transgenic plants, was used as a query to search putative off-target sites across the tomato genome with up to 4 nucleotide mismatches by Geneious R11 or with up to 3 nucleotide mismatches by Cas-OFFinder (Bae *et al*., 2014). Seven potential off-target sites with the highest similarity to the spacer sequence of Wak1-gRNA1 were chosen for evaluation. Genomic regions spanning the putative off-target sites were amplified with specific primers (**Supplemental Table S1**) and PCR amplicons were sequenced to determine if off-target mutations were induced at those sites.

### Bacterial inoculation assay

Four-week-old Δwak1 and wild-type plants were vacuum infiltrated with various *Pst* DC3000 strains at different titers, including DC3000Δ*avrPto*Δ*avrPtoB* (DC3000ΔΔ) or DC3000Δ*avrPto*Δ*avrPtoB*Δ*fliC* (DC3000ΔΔΔ) at 5 x 10^4^ cfu/mL or DC3000 at 1 x 10^6^ cfu/mL. Three to four plants per line were tested with each bacterial strain. Bacterial populations were measured at 3 h and two days after inoculation. Disease symptoms were photographed 4 or 5 days after bacterial infection. Δwak1 and wild-type plants were also spray inoculated with DC3000ΔΔ at 1 x 10^8^ cfu/mL and photographs of disease symptoms were taken 6 days after inoculation.

### PTI protection assay

Four leaflets on the third leaf of 4-week-old plants were first syringe infiltrated with 1 x 10^8^ cfu/mL of heat-killed DC3000Δ*avrPto*Δ*avrPtoB*Δ*hopQ1-1*Δ*fliC* (DC3000ΔΔΔΔ) complemented with a *fliC* allele from DC3000 or ES4326, or no *fliC* (empty vector; EV). Sixteen hours later, the whole plants were vacuum inoculated with DC3000Δ*avrPto*Δ*avrPtoB*Δ*fliC* (DC3000ΔΔΔ) at 5 x 10^4^ cfu/mL. Bacterial populations were measured two days after inoculation. Alternatively, plants were first syringe infiltrated with 1 μM flg22 (GenScript), 1 μM flgII-28 (EZBiolab), or buffer alone (10 mM MgCl_2_), respectively. Plants were inoculated with DC3000ΔΔΔ 16 h later and bacterial populations were measured two days after inoculation as described above.

### Measurement of stomata number and stomata conductance

Leaf samples were taken from Δwak1 and wild-type plants. Photographs from the abaxial epidermis of the leaves were taken using an epifluorescence microscope (Olympus BX51) and the number of cells and both closed and open stomata were counted manually. The stomata index was calculated as the percentage of stomata number per total number of cells (stomata plus epidermal cells). Stomatal conductance was measured at 2 pm, using a leaf porometer (SC1 Decagon Devices, Inc.) on the abaxial side of two leaflets of the third leaf from four plants per line.

### Reactive oxygen species assay

ROS production was measured as described previously (Hind *et al*., 2016). In brief, leaf discs were collected and floated in water overnight (∼16 h). Water was then removed and replaced with a solution containing either 50 nM flg22 (QRLSTGSRINSAKDDAAGLQIA) or 50 nM flgII-28 (ESTNILQRMRELAVQSRNDSNSSTDRDA), in combination with 34 µg/mL luminol (Sigma-Aldrich) and 20 µg/mL horseradish peroxidase. ROS production was then measured over 45 min using a Synergy 2 microplate reader (BioTek). Three to four plants per line and three discs per plant were collected for each experiment.

### Mitogen-activated protein kinase (MAPK) phosphorylation assay

Six leaf discs of Δwak1 and wild-type plants were floated in water overnight to let the wound response subside. The leaf discs were then incubated in 10 nM flg22, 25 nM flgII-28, or water (negative control) for 10 min, and immediately frozen in liquid nitrogen. Protein was extracted using a buffer containing 50 mM Tris-HCl (pH7.5), 10% glycerol, 2 mM EDTA, 1% Triton X-100, 5 mM DTT, 1% protease inhibitor cocktail (Sigma-Aldrich), 0.5% Phosphatase inhibitor cocktail 2 (Sigma-Aldrich). MAPK phosphorylation was determined using an anti-phospho-p44/42 MAPK (Erk1/2) antibody (anti-pMAPK; Cell Signaling).

### Callose deposition

Four-week-old plants were vacuum infiltrated with 1 x 10^8^ cfu/mL *P. fluorescens* 55, a strong inducer of PTI (Rosli *et al*., 2013). Leaf samples were taken 24 h post infiltration, cleared with 96% ethanol and stained with aniline blue for 1 h. Callose deposits were analyzed using an epifluorescence microscope (Olympus BX51). Quantification was performed using ImageJ software. Fifteen photographs per biological replicate were analyzed using four plants per line.

### Co-immunoprecipitation

*Agrobacterium* strains (GV3101+ pMP90) carrying a Gateway binary vector with *Fls2*, *Fls3*, *Bak1*, *Wak1* or *GFP*/*YFP* were infiltrated into leaves of four-week-old *N. benthamiana*. Leaves were treated with either 1 μM flg22, 1 μM flgII-28, or buffer alone for 2 minutes before harvesting. Total protein was extracted from 500 mg *N. benthamiana* leaves in 1.5 mL extraction buffer consisting of 50 mM Tris-HCl (pH 7.5), 150 mM NaCl, 0.5% Triton X-100, 1% (v/v) plant protease inhibitor cocktail (Sigma-Aldrich), 1mM Na_3_VO_4_, 1 mM NaF, and 20 mM β-glycerophosphate. Soluble proteins were incubated with 20 μl of GFP-Trap_MA slurry (Chromotek) or anti-Myc magnetic beads (ThermoFisher Scientific) per sample for 2 h at 4°C, followed by washing three times with cold extraction buffer, and one more wash with cold 50 mM Tris-HCl (pH 7.5). Eluted proteins with 40 μl 2X Laemmli sample buffer and boiled at 95°C for 5 min. For input samples, 8 μL soluble protein mixed with 2X sample buffer were loaded for gel electrophoresis.

### Reverse transcriptase quantitative PCR

Four leaflets from the third leaf of 5-week-old plants were first syringe infiltrated with 1 μM flgII-28 or buffer. Three plants were used for each treatment and two biological replicates were performed. Leaf tissues were collected 0.5 h, 1 h, 2 h, 4 h, 6 h, 8 h after infiltration, immediately frozen in liquid N_2_ and stored at -80°C until used. Total RNA was isolated using RNeasy Plant Mini Kit (Qiagen). RNA (4 μg) was treated with TURBO DNA-free DNase (ThermoFisher Scientific) twice, each for 30 min at 37°C. First-strand cDNA was synthesized from 2 μg RNA using SuperScript^TM^ III (ThermoFisher Scientific). Quantitative PCR was performed with specific primers (**Table S1**) using the QuantStudio™ 6 Flex Real-Time PCR System (ThermoFisher Scientific) and cycling conditions for PCR were 50°C for 2 min, 95°C for 10 min, and 40 cycles of 95°C for 30 s, 56°C for 30 s and 72°C for 30 s.

## Results

### Generation of *Wak1* mutants in tomato by CRISPR/Cas9

We reported previously that virus-induced gene silencing of three homologs of *Wak1* in *N. benthamiana* led to enhanced susceptibility to *Pseudomonas syringae* pv. tomato (Rosli *et al*., 2013). In tomato leaves, transcript abundance of the *Wak1* gene (Solyc09g014720) is significantly increased after treatment with flg22, flgII-28, or csp22, suggesting *Wak1* might play a role in tomato-*Pst* interactions (Rosli *et al*., 2013; Pombo *et al*., 2017). To study the possible role of *Wak1* in plant immunity, we generated mutations in *Wak1* using CRISPR/Cas9 with a guide RNA, Wak1-gRNA1 (GTTAAGATTAGCATAAAACA; **Fig. 1a**), which targets the first exon of the *Wak1* gene. After transformation of the cultivar Rio Grande-PtoR (RG-PtoR, which has the *Pto* and *Prf* genes), we obtained a biallelic mutant (Δwak1 4) from which two *Wak1* homozygous mutant lines (Δwak1 4-1; Δwak1 4-2) were derived (**Fig. 1a**). Line 4-1 has a 10-bp deletion in *Wak1*, resulting in a premature stop codon at the 17^th^ amino acid (aa) of the protein, whereas line 4-2 has a 1-bp deletion in *Wak1*, causing a premature stop codon at the 18^th^ aa (**Fig. 1a**). The growth, development and overall morphology of both Δwak1 mutants were indistinguishable from wild-type RG-PtoR plants (**Fig. S1**).

**Figure 1.**
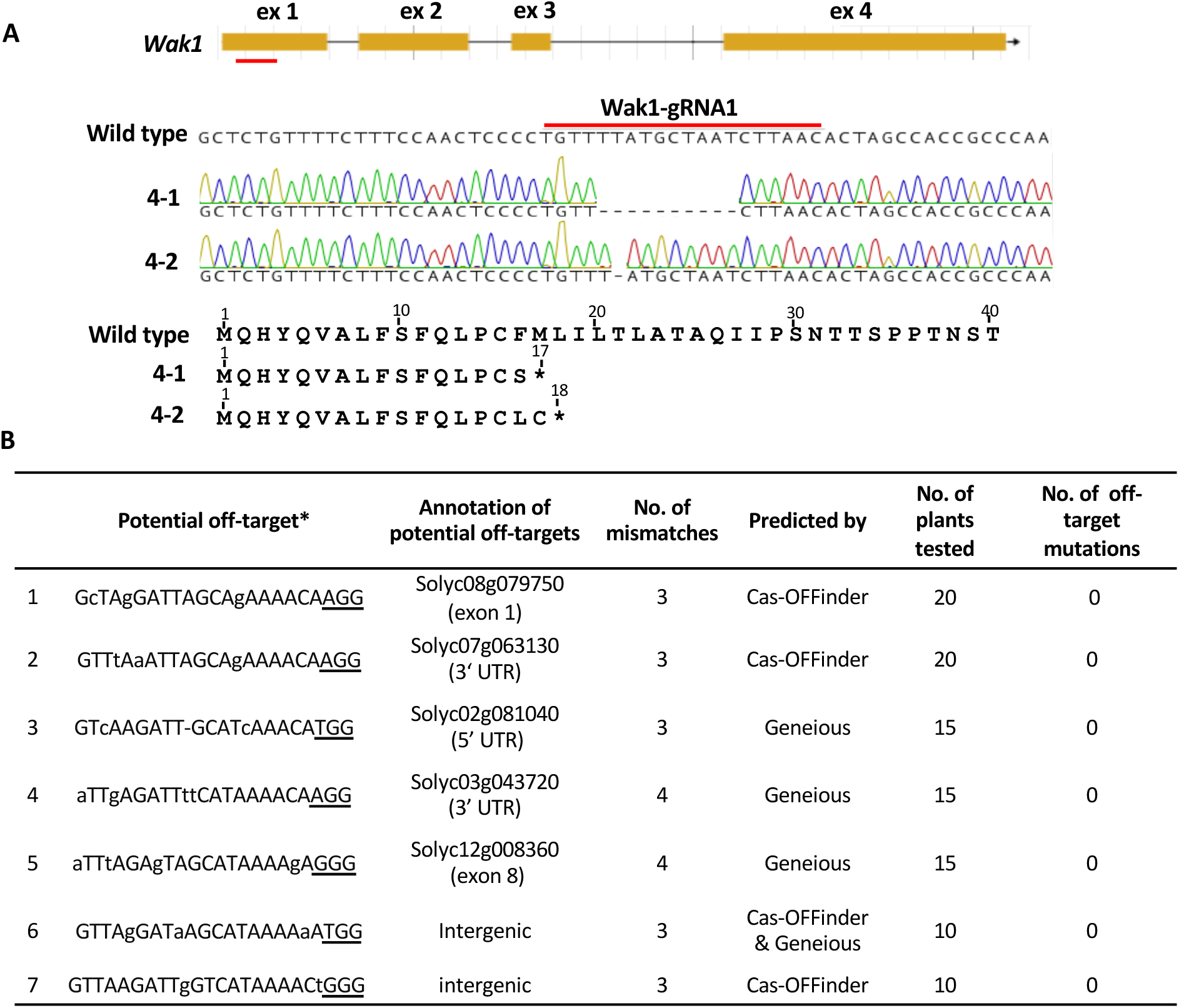
Generation of tomato *Wak1* mutants by CRISPR/Cas9. **A)** Schematics showing the guide-RNA (gRNA) target site in exon 1 (ex 1) and the missense mutations present in two Δwak1 lines (4-1 and 4-2). The gRNA was designed to target the first exon of the *Wak1* gene. The Δwak1 4-1 line has a 10-bp deletion and the Δwak1 4-2 line has a 1-bp deletion. Ex, exon; wild type is RG-PtoR. The Δwak1 lines have a premature stop codon at the 17^th^ or 18^th^ amino acid of the Wak1 protein. **B)** No mutations were detected in any of the potential off-target sites of the Δwak1 plants. For each potential off-target site, 10 to 20 individual plants (T1 or T2 plants) were tested. *PAM (NGG) is underlined; mismatching bases are shown in lowercase.

To determine if the gRNA designed for *Wak1* editing inadvertently caused mutations in other genomic regions of the Δwak1 plants, we selected seven putative sites with the highest off-target scores using Geneious R11 and Cas-OFFinder, although all of these sites had at least three mismatches compared with the spacer sequence of the *Wak1* gRNA (**Fig. 1b**). Of the seven potential off-target sites, two are located in the coding region of a gene, three are in the untranslated region of genes, and another two are in intergenic regions. For each site, we tested 10 to 20 independent T1 or T2 plants, with or without Cas9, and did not detect any off-target mutations. This is not unexpected as the gRNA we designed for *Wak1* was highly specific, with little possibility to target *Wak1* homologs or other genes in tomato, considering that even one mismatch in the seed sequence (the last 12 nucleotides of a gRNA spacer sequence) can severely impair or completely abrogate the editing ability of the Cas9/gRNA complex (Jiang & Doudna, 2017).

### Δwak1 plants are compromised in PRR-triggered, but not NLR-triggered immunity against *Pseudomonas syringae* pv. tomato

To test whether PTI responses are affected in the Δwak1 plants we vacuum-infiltrated Δwak1 and wild-type RG-PtoR plants with the *Pst* strain DC3000Δ*avrPto*Δ*avrPtoB* (DC3000ΔΔ), in which *avrPto* and *avrPtoB* have been deleted and therefore cannot activate NTI. Both Δwak1 lines showed enhanced disease symptoms compared to wild-type plants 4 days after bacterial inoculation, with about six-fold more bacterial growth compared to the wild-type plants (**Fig. 2a**). No differences in symptoms or bacterial populations were observed between the Δwak1 and wild-type plants when they were inoculated with DC3000Δ*avrPto*Δ*avrPtoB*Δ*fliC* (DC3000ΔΔΔ; **Fig. 2b**), which lacks *avrPto* and *avrPtoB* and the flagellin-encoding gene *fliC*. This result indicates that Wak1 plays a role in flagellin-mediated PTI.

**Figure 2.**
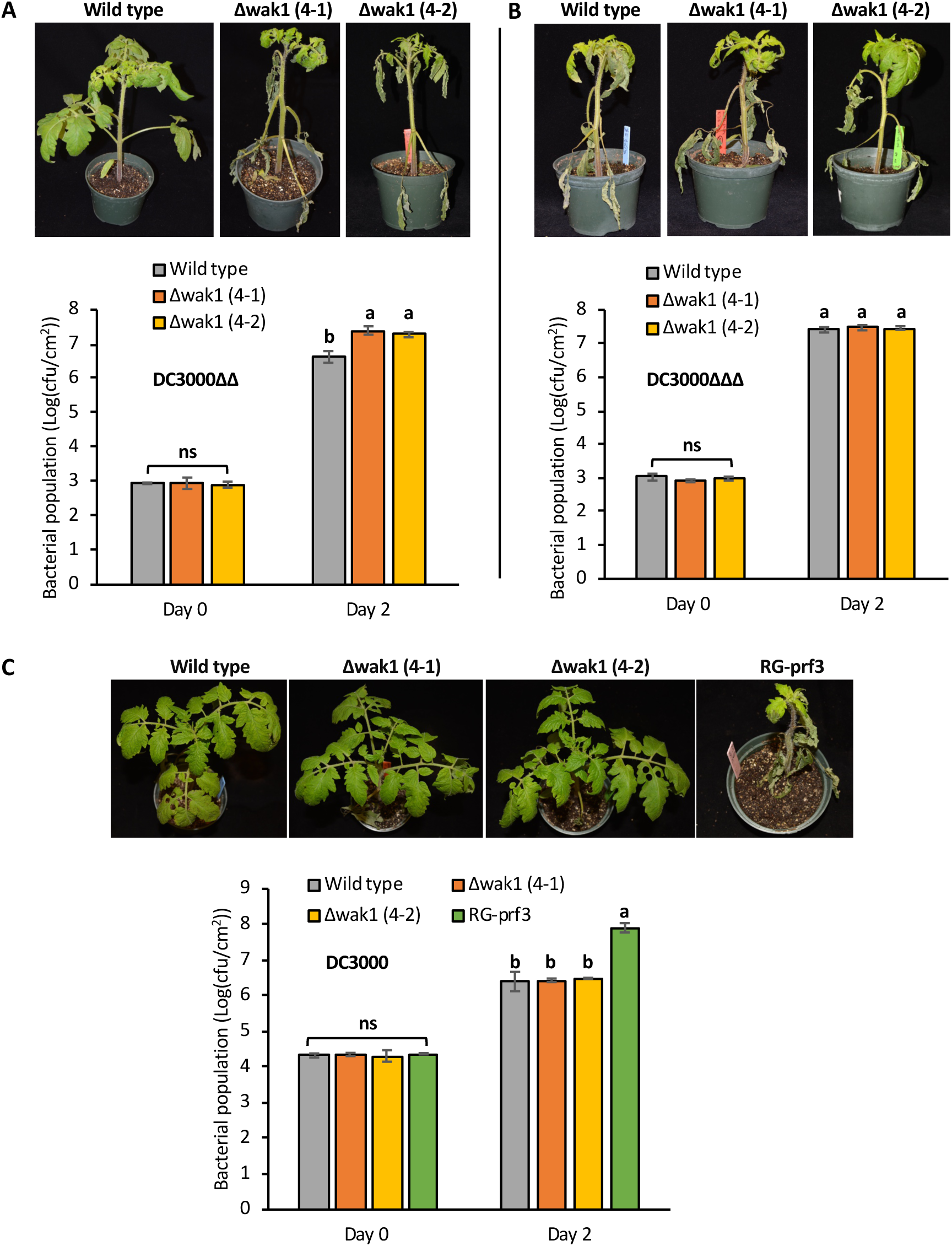
The Δwak1 tomato plants are compromised in flagellin-mediated PRR-triggered immunity but unaffected in NLR-triggered immunity. **A-C)** Four week-old Δwak1 plants and wild-type RG-PtoR plants were vacuum-infiltrated with 5 x 10^4^ cfu/mL DC3000Δ*avrPto*Δ*avrPtoB* (DC3000ΔΔ) (**A**), or 5 x 10^4^ cfu/mL DC3000Δ*avrPto*Δ*avrPtoB*Δ*fliC* (DC3000ΔΔΔ) (**B**), or 1 x 10^6^ cfu/mL DC3000 (C). Photographs of disease symptoms were taken 4 days (**A, B**) or 6 days (**C**) after inoculation. Bacterial populations were measured at 3 hours (Day 0) and two days (Day 2) after infiltration. Bars show means ± standard deviation (SD). Different letters indicate significant differences based on a one-way ANOVA followed by Tukey’s HSD post hoc test (p < 0.05). ns, no significant difference. Three or four plants for each genotype were tested per experiment. The experiment was performed three times with similar results.

To test whether Wak1 contributes to NTI, Δwak1, RG-PtoR, and Rio Grande-prf3 plants (RG-prf3, which contains a mutation in *Prf* that makes the Pto pathway nonfunctional) were inoculated with DC3000. Six days after inoculation, the Δwak1 and RG-PtoR plants had no disease symptoms, whereas the RG-prf3 control showed severe disease symptoms (**Fig. 2c**). Bacterial populations were about 30-fold less in the Δwak1 and RG-PtoR plants compared to RG-prf3. Wak1 therefore appears to have no observable role in NTI.

The two Δwak1 mutant lines were derived from the same primary transformant and it was formally possible that another mutation induced during tissue culture is responsible for the enhanced susceptibility to *Pst*. We therefore developed F1 hybrids by crossing the Δwak1 plants to RG-PtoR plants (**Fig. S2**). Sequencing confirmed that all F1 hybrids were heterozygous for the *Wak1* mutation. F1 hybrids that were vacuum-infiltrated with DC3000ΔΔ developed disease symptoms and supported bacterial populations similar to RG-PtoR plants (**Fig. S2a**), indicating *Wak1* is a dominant allele. Four F1 plants (two were -10 bp/WT and two were -1 bp/WT; WT, wild type) were selfed to develop F2 populations. After inoculation of 117 F2 plants with DC3000ΔΔ we observed a segregation ratio of 3 resistant to 1 susceptible (**Fig. S2b**). Sequencing revealed all resistant plants were either homozygous wildtype or heterozygous, while the susceptible plants were homozygous for the *wak1* mutation (**Fig. S2c**). Combined with the lack of off-target mutations, these disease assays with F2 populations strongly support that the susceptibility to *Pst* of Δwak1 plants is due to the CRISPR/Cas9-induced loss-of-function mutations in the *Wak1* gene.

### *Wak1* mutant plants are compromised in PRR-triggered immunity induced by flg22 and flgII-28

The observation that Δwak1 plants are more susceptible to DC3000ΔΔ but show no differences compared to wild-type plants for their response to DC3000ΔΔΔ which lacks flagellin, suggests that Wak1 is involved in immune responses mediated by flg22 and/or flgII-28. To further test this, we performed a ‘PTI protection’ assay using a heat-killed *Pst* strain lacking flagellin and three type III effectors (DC3000Δ*avrPto*Δ*avrPtoB*Δ*hopQ1-1*Δ*fliC*; DC3000ΔΔΔΔ) complemented with a construct expressing *fliC* from either DC3000 (which has active flg22 and flgII-28) or *P. cannabina* pv. alisalensis ES4326 (only flgII-28 is active) (Hind *et al*., 2016), or an empty vector (EV) as a control (**Fig. 3a**). Since both of the Δwak1 lines were similarly susceptible to DC3000ΔΔ, most subsequent experiments were focused on the 4-1 line. Δwak1 4-1 plants were first infiltrated with the various suspensions of heat-killed bacteria to induce PTI and then challenged with DC3000ΔΔΔ 16 h later. Wild-type plants pretreated with *Pst* DC3000ΔΔΔΔ with an empty vector supported a significantly higher bacterial population than plants pretreated with the heat-killed bacterial suspensions containing either DC3000 *fliC* or ES4326 *fliC* (7.5-fold and 3.3-fold, respectively), indicating that pretreatment of wild-type plants activated PTI defenses due to recognition of flg22 and/or flgII-28. The Δwak1 plants, however, supported higher bacterial populations regardless of the pretreatment indicating the PTI response was compromised (**Fig. 3a**).

**Figure 3.**
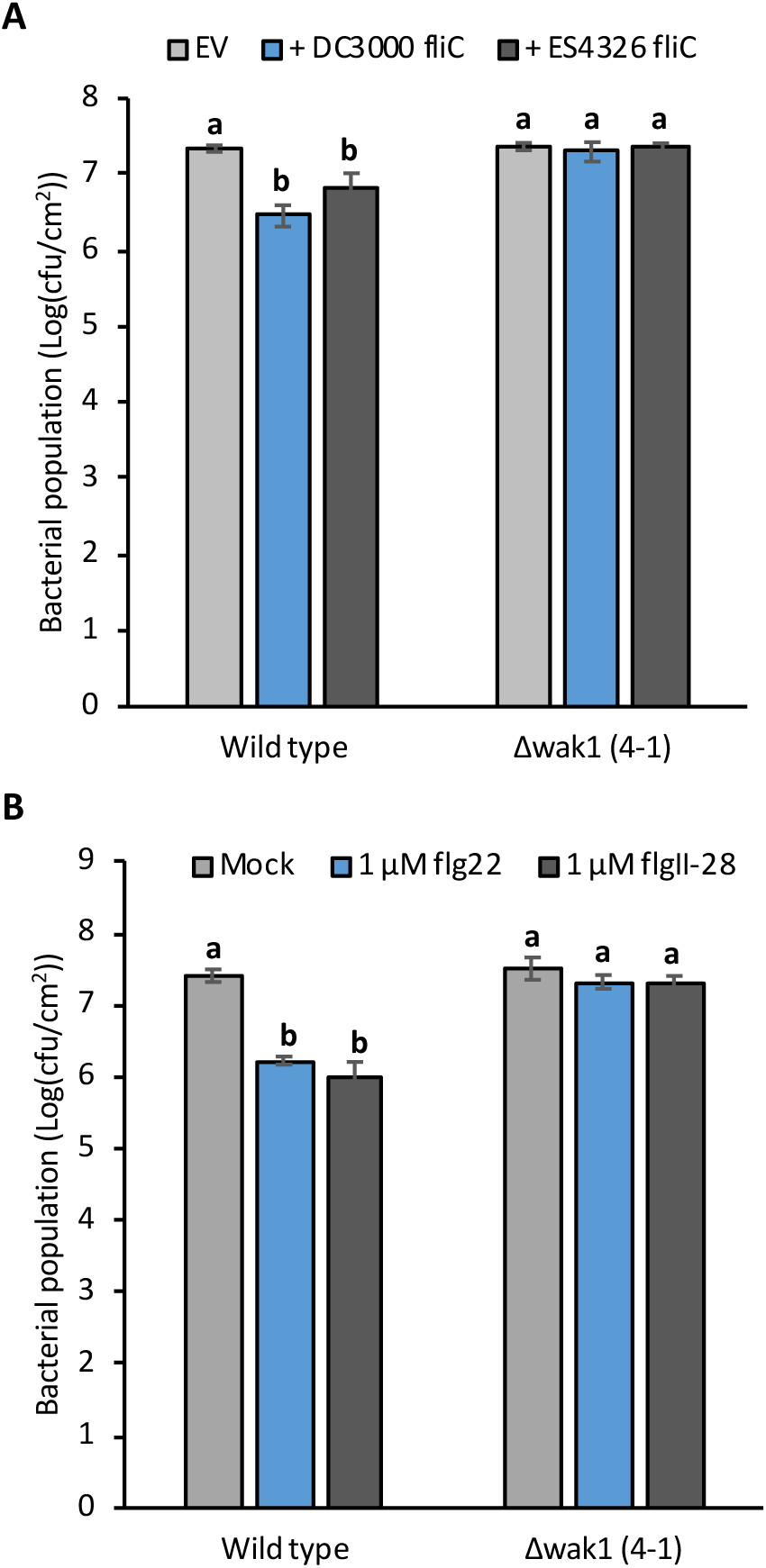
The Δwak1 plants are compromised in two PRR-triggered immunity induction assays. **A)** Four week-old Δwak1 plants (4-1) and wild-type RG-PtoR plants were first syringe infiltrated with 1 x 10^8^ cfu/mL of heat-killed DC3000Δ*avrPto*Δ*avrPtoB*Δ*hopQ1-1*Δ*fliC* (DC3000ΔΔΔΔ) complemented with a *fliC* gene from DC3000 or ES4326, or no *fliC* (empty vector, EV). Sixteen hours later, the whole plants were vacuum infiltrated with DC3000Δ*avrPto*Δ*avrPto*BΔ*fliC* (DC3000ΔΔΔ) at 5 x 10^4^ cfu/mL. Bacterial populations were measured two days after the infiltration. **B**) Plants (Δwak1 4-1 and wild type) were first syringe infiltrated with buffer only (mock; 10 mM MgCl_2_), 1 μM flg22, or 1 μM flgII-28, respectively. Sixteen hours later, plants were vacuum infiltrated with DC3000ΔΔΔ at 5 x 10^4^ cfu/mL. Bacterial populations were measured two days later. Bars in (**a**) and (**b**) represent means ± SD. Different letters indicate significant differences based on a one-way ANOVA followed by Tukey’s HSD post hoc test (p < 0.05).

We next performed the PTI protection assay using the synthetic peptides flg22 and flgII-28. Plants were first syringe-infiltrated with buffer alone, 1 μM flg22, or 1 μM flgII-28, and then challenged with DC3000ΔΔΔ 16 h later as described above (**Fig. 3b**). Two days later, wild-type plants that were pretreated with either flg22 or flgII-28 had significantly lower bacterial populations compared to the buffer-only treatment. In contrast, no significant differences in bacterial populations regardless of pretreatment were observed in Δwak1 plants. Collectively, these experiments demonstrate that Wak1 plays an important role in PTI that is activated by two flagellin-derived MAMPs.

### Δwak1 plants are not compromised in PRR-triggered immunity responses on the leaf surface, or in stomatal numbers or conductance

*Pst* inoculation experiments using vacuum infiltration assess PTI responses primarily in the apoplast. To test if Wak1-mediated immunity also plays a role in PTI on the leaf surface, we spray inoculated Δwak1 and wild-type RG-PtoR plants with DC3000ΔΔ. This inoculation method requires the pathogen to enter the apoplastic space through stomata or natural openings. Interestingly, in contrast to experiments using vacuum infiltration, both wild-type and Δwak1 plants developed disease symptoms after spray inoculation that were indistinguishable both in the amount of time until they developed and in their ultimate severity (**Fig. 4a**). Thus, Wak1 does not appear to play an important role in PTI responses on the leaf surface. Measurements of stomatal numbers and of stomatal conductance as an indicator of stomatal activity revealed no differences between wild-type and Δwak1 plants, further indicating that Wak1 does not play a role at the leaf surface (**Figs. 4b,c**).

**Figure 4.**
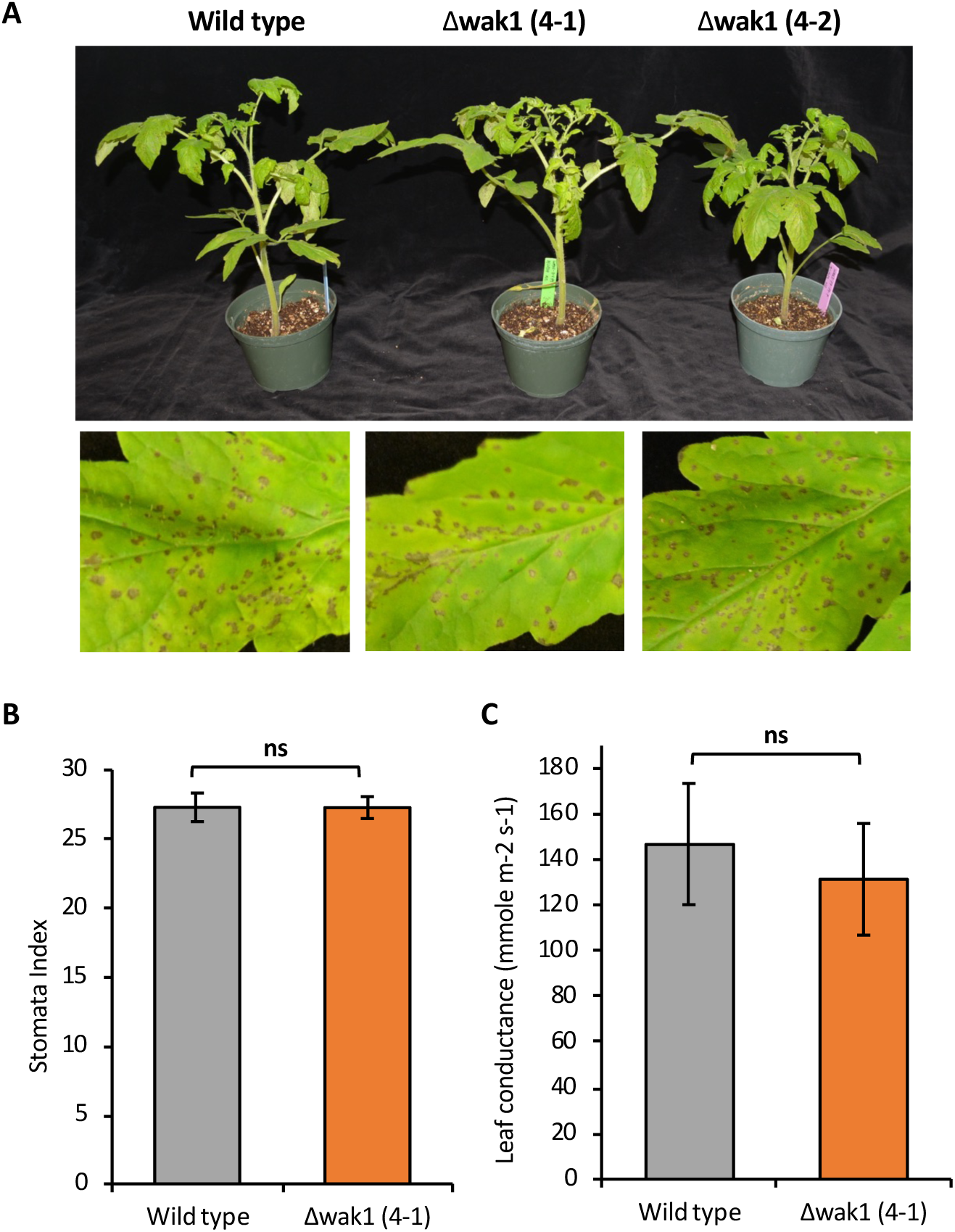
Leaf surface-associated immune responses and stomata are unaffected in Δwak1 plants. **A**) Four week-old Dwak1 plants and wild-type RG-PtoR plants were spray inoculated with 1 x 10^8^ cfu/mL DC3000Δ*avrPto*Δ*avrPtoB*. Photographs of disease symptoms were taken 6 days after inoculation. Photographs show a representative plant and leaflet from each line. **B**) Stomatal index taken from wild-type and Δwak1 4-1 plants. Photographs from the abaxial epidermis of the leaves were taken using an epifluorescence microscope and the number of cells and both closed and open stomata were counted manually. The stomatal index was calculated as the percentage of stomata number per total number of cells (stomata plus epidermal cells). Five photographs per biological replicate were analyzed. Bars represent the mean of 4 biological replicates with their corresponding standard deviation. **C**) Stomatal conductance measured on the abaxial side of leaflets on the third leaf. Data correspond to the average of two leaflets from at least 4 biological replicates per line, with ± SD. ns, no significant difference using Student’s *t*-test (*p* <0.05).

### Δwak1 plants are unaffected in MAMP-induced ROS production or MAPK activation but have significantly reduced callose deposition

Generation of reactive oxygen species (ROS) and activation of mitogen-activated protein kinase (MAPK) cascades are two typical PTI-associated responses in plants (Nguyen *et al*., 2010; Zipfel, 2014). To investigate whether Wak1 participates in these responses we performed ROS assays and MAPK activation assays using flg22 or flgII-28. We observed no differences in ROS production in Δwak1 plants compared to wild-type plants when treated with either of these flagellin-derived MAMPs (**Fig. 5a,b**). Similarly, we observed no difference between wild-type and Δwak1 plants for their ability to activate MAPKs in response to these two MAMPs (**Fig. 5c**).

**Figure 5.**
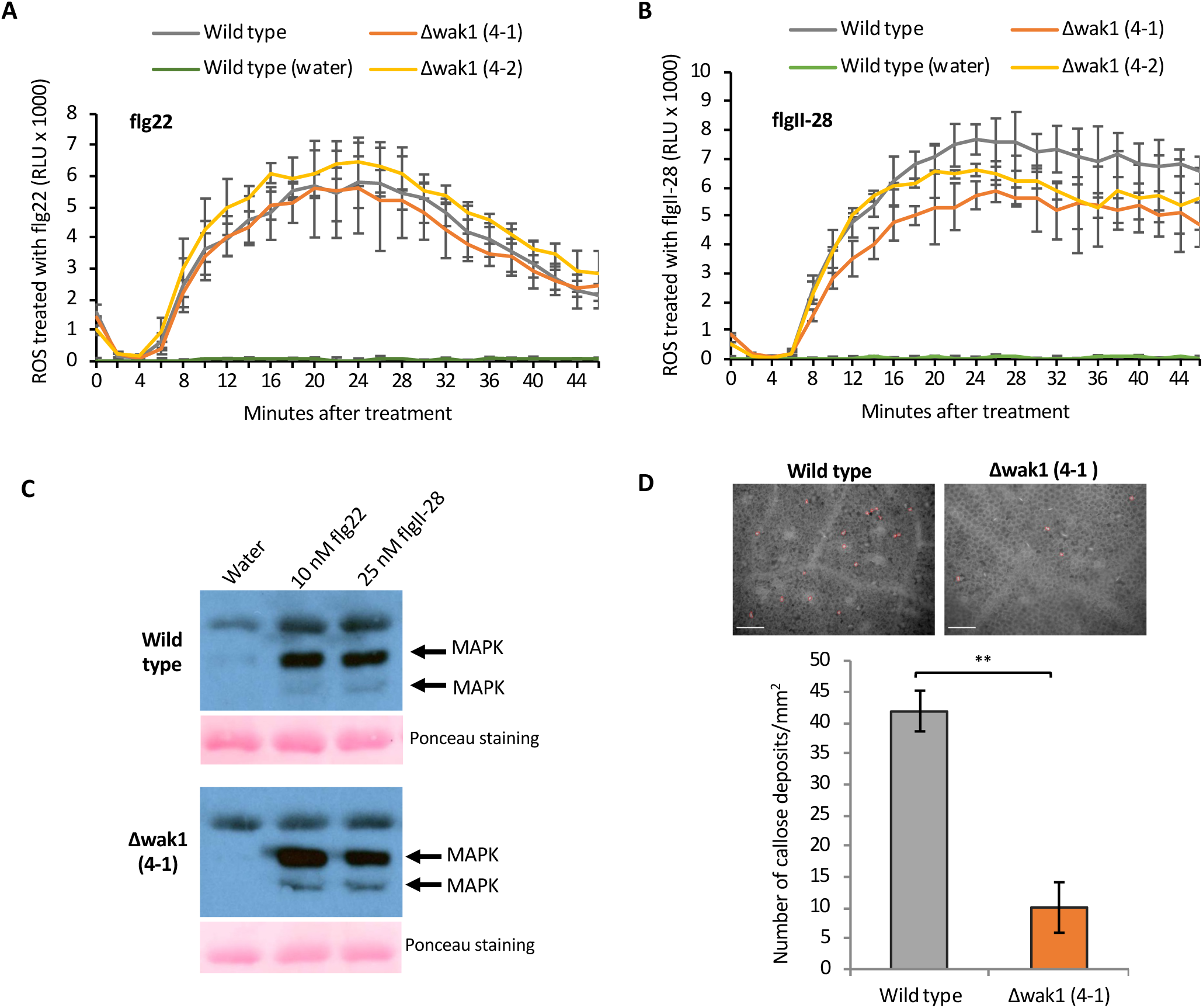
The Δwak1 plants are not affected in MAMP-induced ROS production or MAPK activation but have reduced callose deposition. (**A-B**) Leaf discs from Δwak1 or wild-type plants were treated with 50 nM flg22 (**A**), 50 nM flgII-28 (**B**), or water only, and relative light units (RLU) were measured over 45 minutes. One-way ANOVA followed by Tukey’s HSD post hoc test (p < 0.05) was performed at 24 min (peak readout) and 45 min after treatment with flg22 or flgII-28. No significant difference was observed between Δwak1 and wild-type plants in either treatment. **C**) Leaf discs from Δwak1 (4-1) or wild-type RG-PtoR plants were treated with water, 10 nM flg22, or 25 nM flgII-28 for 10 min. Proteins were extracted from a pool of discs from three plants and subjected to immunoblotting using an anti-pMAPK antibody that detects phosphorylated MAPKs. Ponceau staining shows equal loading of protein. This experiment was performed three times with similar results. D) Wild-type and Δwak1 plants (4-1) were vacuum infiltrated with 1 x 10^8^ cfu/mL *Pseudomonas fluorescens* 55. Leaf samples were taken 24 h after infiltration, de-stained with 96% ethanol and stained with aniline blue for 1 h. Callose deposits were analyzed using an epifluorescence microscope. Top: Representative photographs of wildtype and Δwak1 plants taken for callose deposition estimation. Red spots indicate the callose deposits observed and used for quantification. Scale bars: 100 μm. Bottom: Total number of callose deposits per mm^2^ quantified in each group of plants. Fifteen photographs per biological replicate were analyzed. Bars represent the mean of 4 biological replicates with their corresponding standard deviation. The asterisks represent a significant difference using Student’s *t*-test (*p* <0.01).

Callose deposition is a response associated with later stages of PTI, and one which is regulated independently or downstream of MAPK activation (He et al., 2016). We measured callose deposition by challenging Δwak1 and wild-type plants using a non-pathogenic bacterial strain, *P. fluorescens* 55, a strong inducer of PTI (Rosli *et al*., 2013). Compared to wild-type plants, Δwak1 plants showed significantly reduced callose deposition one day after vacuum infiltration of *Pf* 55 (**Fig. 5d**). These observations therefore indicate that Wak1 functions at a later stage of the PTI response in a flagellin-induced signaling pathway independent of ROS production and MAPK activation.

### The increase in *Wak1* transcript abundance upon flgII-28 treatment is Fls3-dependent

In tomato, the transcript abundance of *Wak1* is low in unchallenged conditions, but is significantly higher after *Pst* inoculation (Rosli *et al*., 2013). To gain insight into the transcriptional regulation of *Wak1* and *Fls3* during the immune response, we used RT-qPCR to measure *Wak1* and *Fls3* transcript abundance after treatment of wild-type leaves with flgII-28 (**Fig. 6a**). The relative abundance of *Wak1* or *Fls3* transcripts at various time points after syringe infiltrating 1μM flgII-28 was compared to a mock treatment (10 mM MgCl_2_). Both *Wak1* and *Fls3* transcript abundance increased significantly at 6 and 8 hours after syringe infiltrating flgII-28 compared to the mock control (**Fig. 6a**).

**Figure 6.**
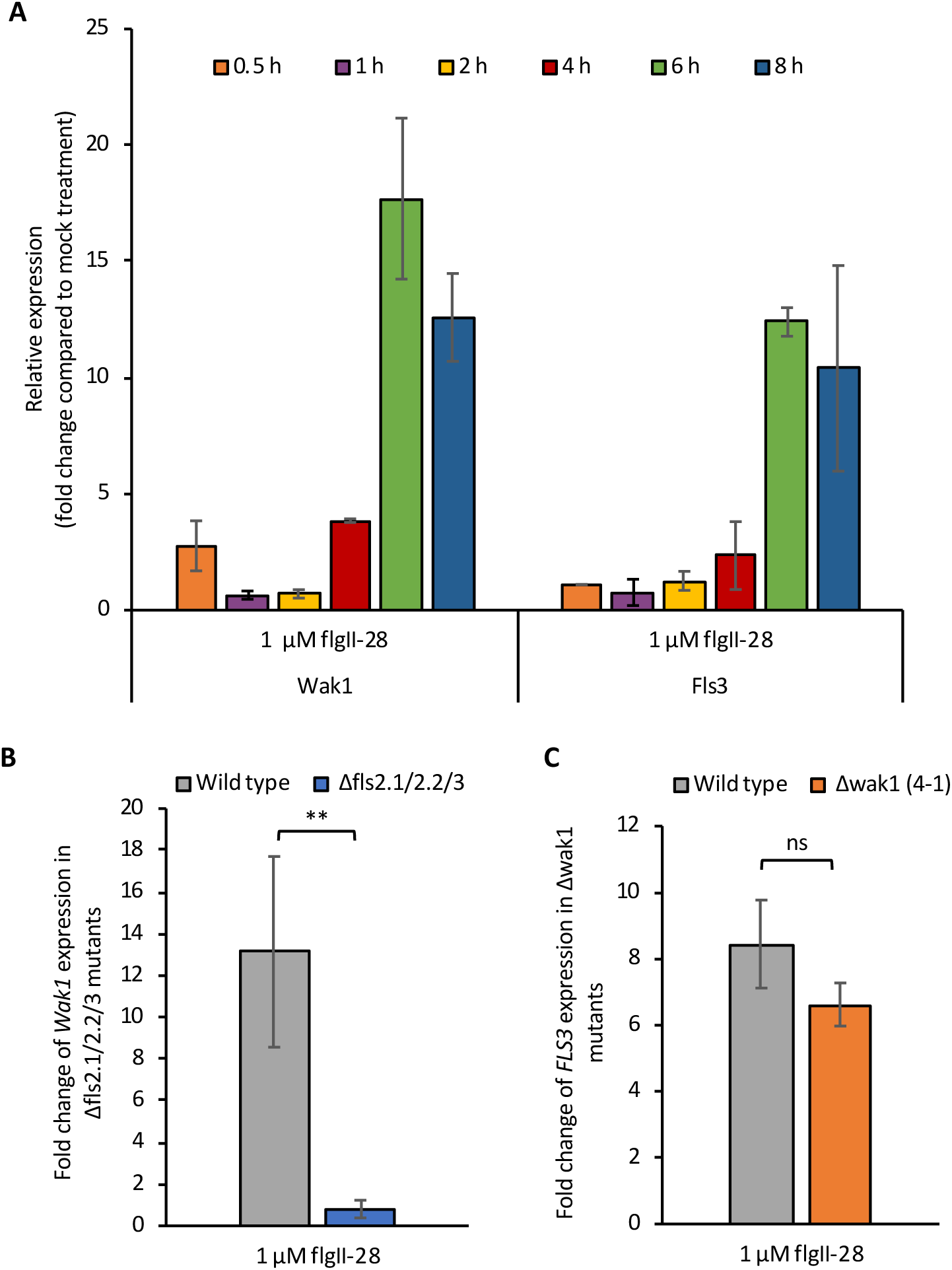
Transcript abundance changes of *Wak1* are dependent on the presence of Fls3 in tomato. **A**) Transcript abundance of *Wak1* and *Fls3* genes measured by RT-qPCR at the times shown after treatment with 1 µM flgII-28 compared to a buffer-only control (10 mM MgCl_2_; mock treatment). Each treatment included three biological replicates and three technical replicates. *SlArd2* (Solyc01g104170) was used as the reference gene for quantification. Bars represent means ± SD. **B**) RT-qPCR was used to measure transcript abundance of *Wak1* 6 h after treatment of Δfls2.1/2.2/3 or wild-type leaves with 1 µM flgII-28. Bars represent the mean ± SD. Two asterisks represent a significant difference using Student’s *t*-test (*p* <0.01). **C**) RT-qPCR was used to measure transcript abundance of *Fls3* 6 h after treatment of Δwak1 (4-1) or wild-type leaves with 1 µM flgII-28. Bars represent the mean ± SD. ns, no significant difference using Student’s *t*-test (*p* <0.05).

To investigate possible co-dependence of *Wak1* and *Fls3* gene expression, we measured the *Wak1* transcript abundance in tomato plants that have mutations in the two *Fls2* genes and *Fls3* (Δfls2.1/2.2/3; (Roberts *et al*., 2020)) and the *Fls3* transcript abundance in Δwak1 plants after treatment with flgII-28. The abundance of *Wak1* transcripts was greatly reduced in the Δfls2.1/2.2/3 plants compared to wild-type plants, whereas *Fls3* abundance was not significantly different in Δwak1 or wild-type plants (**Fig. 6b,c**). These results indicate that *Wak1* gene expression is regulated by the Fls3 pathway and its function likely occurs downstream of the mechanism inducing *Fls3* gene expression.

### Δwak1 plants develop bacterial speck disease symptoms more slowly than Δfls2.1/2.2/3 plants

To determine the relative contributions of Wak1 and Fls2/Fls3 to PTI we next compared the response of Δwak1 and Δfls2.1/2.2/3 plants to DC3000ΔΔ (**Fig. 7**). Three days after inoculation, the Δfls2.1/2.2/3 plants showed more severe disease symptoms than Δwak1 plants or wild-type plants, but by 4 days after inoculation both the Δwak1 and Δfls2.1/2.2/3 plants developed more disease symptoms than the wild-type plants (**Fig. 8a**). There was no distinguishable difference between the Δwak1 and Δfls2.1/2.2/3 plants 4-10 days after inoculation (**Fig. 8a**). Two days after inoculation, the bacterial population in the Δfls2.1/2.2/3 and Δwak1 plants was 6-fold and 4-fold higher than the wild-type plants, respectively, with no statistically significant difference in bacterial populations between the Δwak1 and Δfls2.1/2.2/3 plants (**Fig. 8b**).

**Figure 7.**
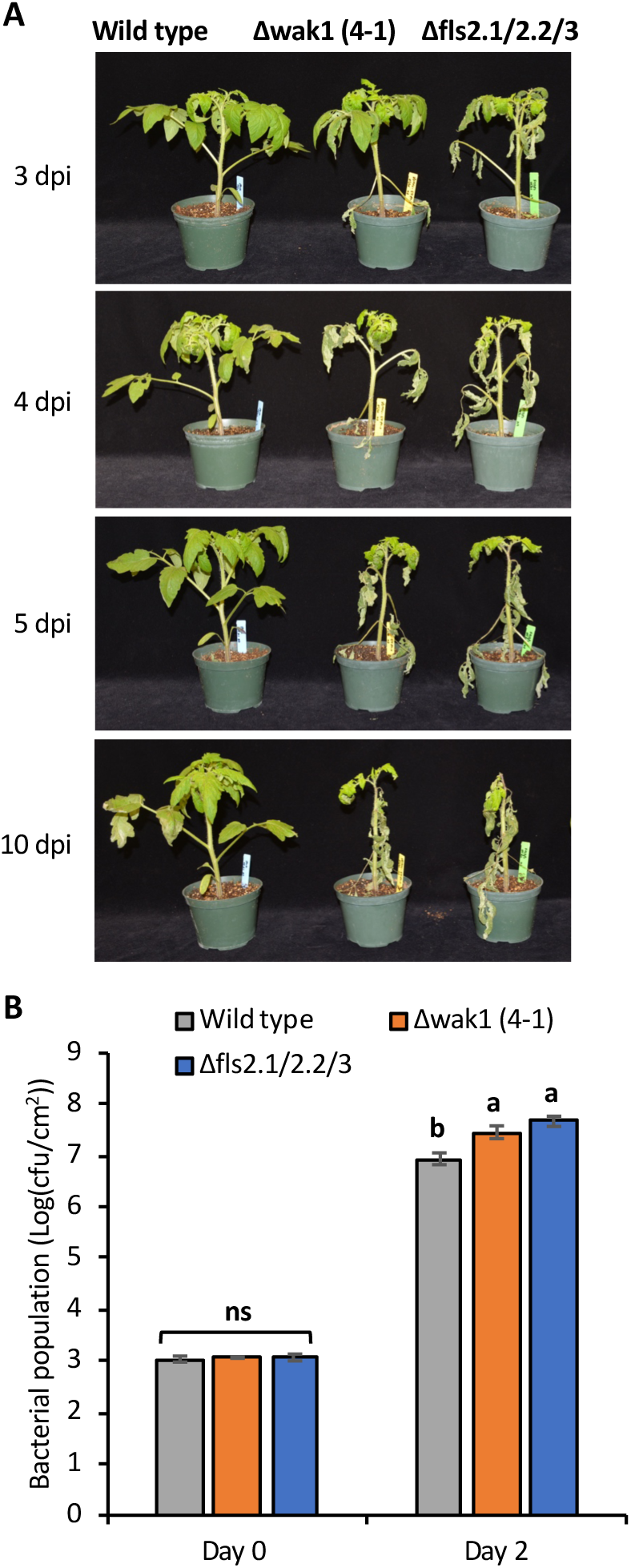
The Δwak1 plants develop disease symptoms more slowly than Δfls2.1/2.2/3 plants. Four week-old Δwak1 (4-1), Δfls2.1/2.2/3 or wild-type RG-PtoR plants were vacuum infiltrated with 5 x 10^4^ cfu/mL DC3000Δ*avrPto*Δ*avrPtoB*. **A**) Photographs were taken at 3, 4, 5, or 10 days after inoculation. **B**) Bacterial populations were measured 3 hours (Day 0) and two days after infiltration (Day 2). Bars represent means ± SD. Different letters indicate significant differences based on a one-way ANOVA followed by Tukey’s HSD post hoc test (p < 0.05). ns, no significant difference. Three or four plants for each genotype were tested per experiment. This experiment was performed twice with similar results

**Figure 8.**
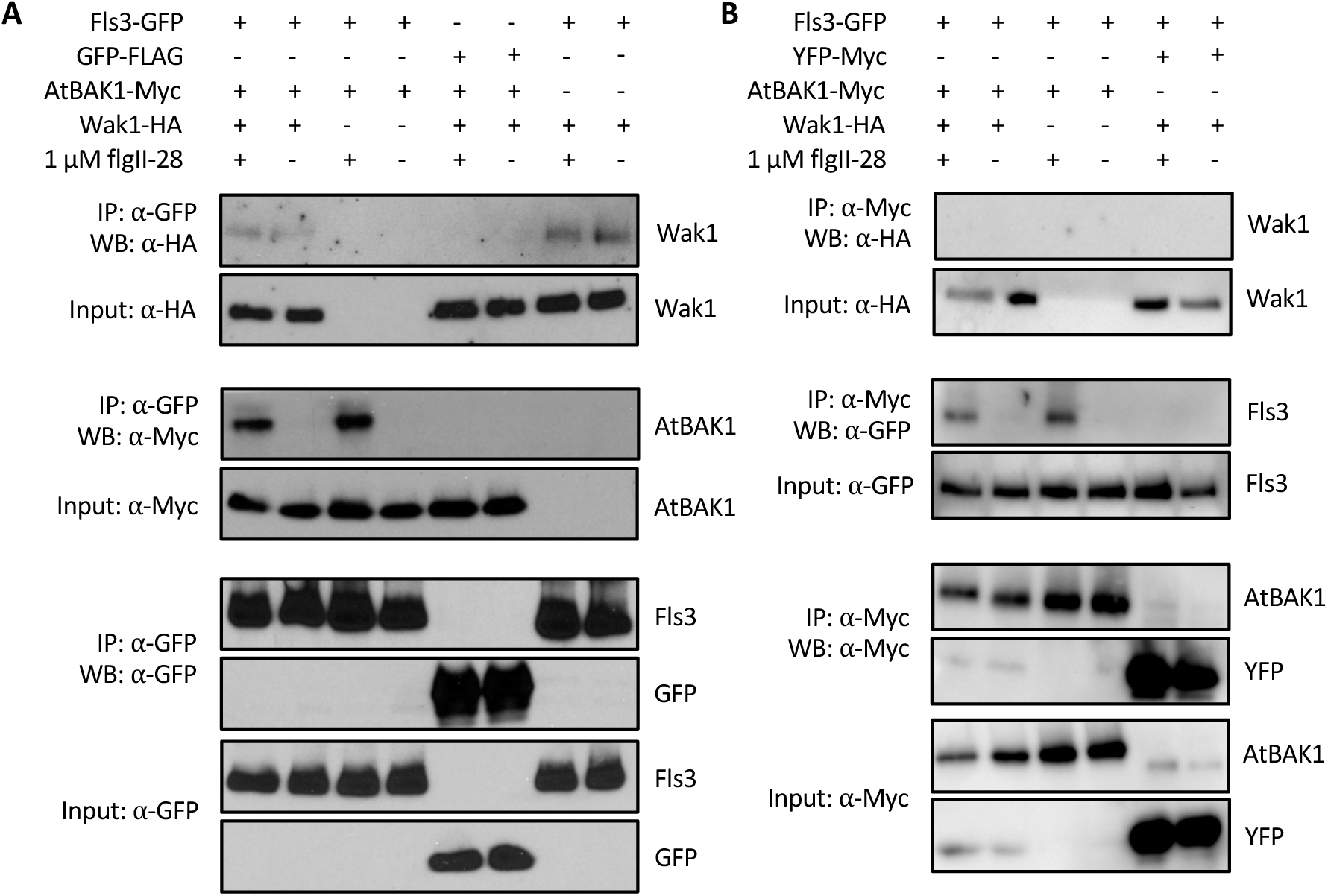
Wak1 associates with Fls3 independently of flgII-28 and BAK1 and Wak1 does not associate with BAK1. **(A-B)** Proteins were extracted from *N. benthamiana* leaves expressing Fls3-GFP in combination with AtBAK1-Myc and/or Wak1-HA after treatment with or without 1 μM flgII-28 for 2 min and were used for immunoprecipitation using anti GFP magnetic agarose beads (A) or anti-Myc magnetic beads (**B**). Wak1 was pulled down with Fls3 (**A**) but not with BAK1 (B) after treatment with or without flgII-28. Wak1, BAK1, Fls3, GFP and YFP proteins were detected by immunoblotting with ⍺-HA, ⍺-Myc, or ⍺-GFP antibodies. This experiment was repeated three times with similar results.

### *Wak1* occurs in a complex with Fls2 and Fls3 independent of flg22, flgII-28 or Bak1

The results above indicate that Wak1 plays a major role in flg22- and flgII-28-induced processes that occur in the apoplast later in the PTI response. We considered the possibility that Wak1 acts in a complex with Fls2 and Fls3 similar to what has been reported for FLS2 and FERONIA in Arabidopsis (Stegmann *et al*., 2017). We therefore used transient expression of proteins in *N. benthamiana* leaves and co-immunoprecipitation (Co-IP) to investigate if Wak1 physically associates with Fls2, Fls3, or the co-receptor Bak1 and, if so, whether the interaction is affected by the presence of flg22 or flgII-28. We observed a weak, but reproducible and specific, interaction of Wak1 with both Fls2 and Fls3 with the interactions occurring independently of flg22, flgII-28, or the presence of Bak1(**Fig. 8a** and **Fig. S3**). As expected, Fls3 and Fls2 each interacted strongly with Bak1 only in the presence of flgII-28 or flg22, respectively. No interaction was observed between Wak1 and Bak1 proteins (**Fig. 8b**). Additionally, Wak1 did not affect the accumulation of the Fls2, Fls3 or Bak1 proteins or vice versa (**Fig. 8** and **Fig. S3**).

## Discussion

The tomato *Wak1* gene was first identified as a *FIRE* gene (flagellin-induced, repressed by effectors) in the immune response against *Pseudomonas syringe* (Rosli *et al*., 2013). When its expression was knocked down by virus-induced gene silencing (VIGS) in *N. benthamiana* the morphology of the plants was unaffected but their ability to activate PTI was compromised leading to more severe disease symptoms and enhanced growth of a virulent *Pst* strain (Rosli *et al*., 2013). The interpretation of these experiments was limited somewhat by the fact that three *N. benthamiana Wak1* homologs were silenced by the tomato *Wak1* VIGS construct and, as is typical for VIGS, their transcripts were not completely eliminated (they were reduced by ∼50%). Thus, whether one, or more, of the *Wak1* homologs in *N. benthamiana* play a role in PTI was unclear as was the degree to which a complete knockout of the *Wak1* genes might affect PTI or affect plant morphology. Here we have addressed these limitations by developing two CRISPR/Cas9-mediated *Wak1* mutants in tomato and used them to investigate the contributions of Wak1 to several PTI-associated responses and to resistance to *P. syringae*. As elaborated upon below, our results indicate *Wak1* gene expression is induced by the Fls2 and Fls3 pathways in tomato, the Wak1 protein associates in a complex with Fls2 and Fls3, and Wak1 plays an important role in later stages of flagellin-induced PTI.

Consistent with our earlier observations of *Wak1*-silenced *N. benthamiana* plants, the Δwak1 tomato plants developed more severe disease symptoms compared to wild-type plants and supported larger populations of *Pst*; they also had wild-type morphology. Interestingly, the differences in pathogen responses were abolished when the *Pst* strain used for inoculation lacked flagellin suggesting that either flg22 and/or flgII-28 and their corresponding receptors Fls2 and Fls3 play a key role in activating Wak1-mediated responses. In fact, subsequent experiments using *Pst* strains with variant FliC proteins or using synthetic flg22 and flgII-28 peptides confirmed that either one of these MAMPs is sufficient to induce Wak1-dependent PTI. At this stage of the work this dependence could be potentially explained simply by the fact that both of these MAMPs are able to significantly up-regulate expression of the *Wak1* gene.

Several observations support the hypothesis that Wak1 acts at a later stage of the PTI response in tomato. First, the Δwak1 plants showed no difference from wild-type plants when *Pst* was spray-inoculated, a method that assays for PTI responses at the leaf surface. The importance of PTI on the leaf surface has been extensively documented in Arabidopsis where a major regulator of this response is the activity of Fls2 in the stomata (Melotto *et al*., 2006; Melotto *et al*., 2008; Melotto *et al*., 2017). Our observations suggest that Wak1 does not act in PTI on the leaf surface but instead exerts its function at a later stage, after *Pst* enters the apoplastic space as simulated by vacuum infiltration. Second, Δwak1 plants showed no defects in their ability to produce ROS or activate MAPKs in response to flg22 and flgII-28. Both of these responses occur early (within minutes) in leaves that are exposed to MAMPs. Third, *Fls3* gene expression induced by flgII-28 was the same in Δwak1 plants as it was in wild-type plants. Transcriptional changes also occur rapidly (within 1 hour) of MAMP treatment (Pombo *et al*., 2017). As expected, the induction of *Wak1* gene expression by flgII-28 was compromised in Δfls2.1/2.2/3 plants. Fourth, the Dwak1 plants produced just 25% of the callose deposits observed in wild-type plants in response to *P. fluorescens*, a source of flagellin and other MAMPs. Callose deposition occurs later than ROS production and MAPK activation and contributes to cell wall strengthening which may inhibit the infection process (Nguyen *et al*., 2010; Voigt, 2014). Finally, the Δwak1 plants developed disease symptoms more slowly than did Δfls2.1/2.2/3 plants. This would be expected if the Δfls2.1/2.2/3 mutations result in the loss of both early (e.g., ROS, MAPK activation, transcriptional reprogramming) and later-stage PTI (callose deposition) whereas the *Wak1* mutation compromises primarily later-stage PTI responses. Importantly, however, both Δwak1 and Δfls2.1/fls2.2/fls3 plants ultimately developed the same severe disease symptoms which demonstrates the critical role that Wak1 plays in the host response to *Pst*.

The dependence of Wak1-mediated PTI on Fls2 and Fls3 activity could be explained, in part, by the induction of *Wak1* gene expression by the Fls2 and Fls3 pathways. However, our observations also raised the possibility that Wak1 resides in a complex that contains Fls2 and Fls3 and its function involves these receptors. We tested this hypothesis and found that Wak1 does co-immunoprecipitate with Fls2 and Fls3 in a MAMP-independent manner and it does not affect accumulation of Fls2/Fls3 proteins. This is reminiscent of the Arabidopsis malectin-like receptor kinase, FERONIA (FER), which was found to weakly associate with Fls2 independent of flg22 treatment and also had no effect on Fls2 accumulation (Stegmann *et al*., 2017). It is possible that Wak1, like FER, may act as an important cell wall-associated scaffold to regulate immune receptor-complex formation. Tomato Wak1 did not co-immunoprecipitate with Bak1, and Bak1 was not required for the Wak1-Fls2/3 interactions. In contrast, FER weakly associates with Bak1 and the interaction is enhanced upon flg22 treatment, but whether Bak1 is required for the weak association of FER-Fls2 was not investigated (Stegmann *et al*., 2017).

Based on our observations, we propose a model for the role of Wak1 in PTI (**Fig. 9**). In this model, *Wak1* transcript abundance is greatly increased upon activation of the PRRs Fls2 and Fls3. We hypothesize this gene expression occurs primarily when *Pst* enters the apoplastic space and that *Wak1* is not expressed in leaf surface or stomatal cells. Increased transcript abundance leads to increased Wak1 protein accumulation and subsequent localization to a cell wall-associated protein complex that contains Fls2 and Fls3 and possibly other PRRs. Wak1 might act as a receptor of a damage-associated molecular pattern (DAMP), such as oligogalacturonides. Binding of such a DAMP might impact the association of Wak1 with the Fls2/Fls3 complex to promote stabilization and accumulation of the PRRs, enhance the interaction of Wak1 with PRRs, or possibly stimulate PRR kinase activity. Whatever the mechanism, the presence of Wak1 in this wall-associated kinase plays a critical role in later stages of PTI including callose deposition and other processes that ultimately inhibit growth of virulent *Pst*.

**Figure 9.**
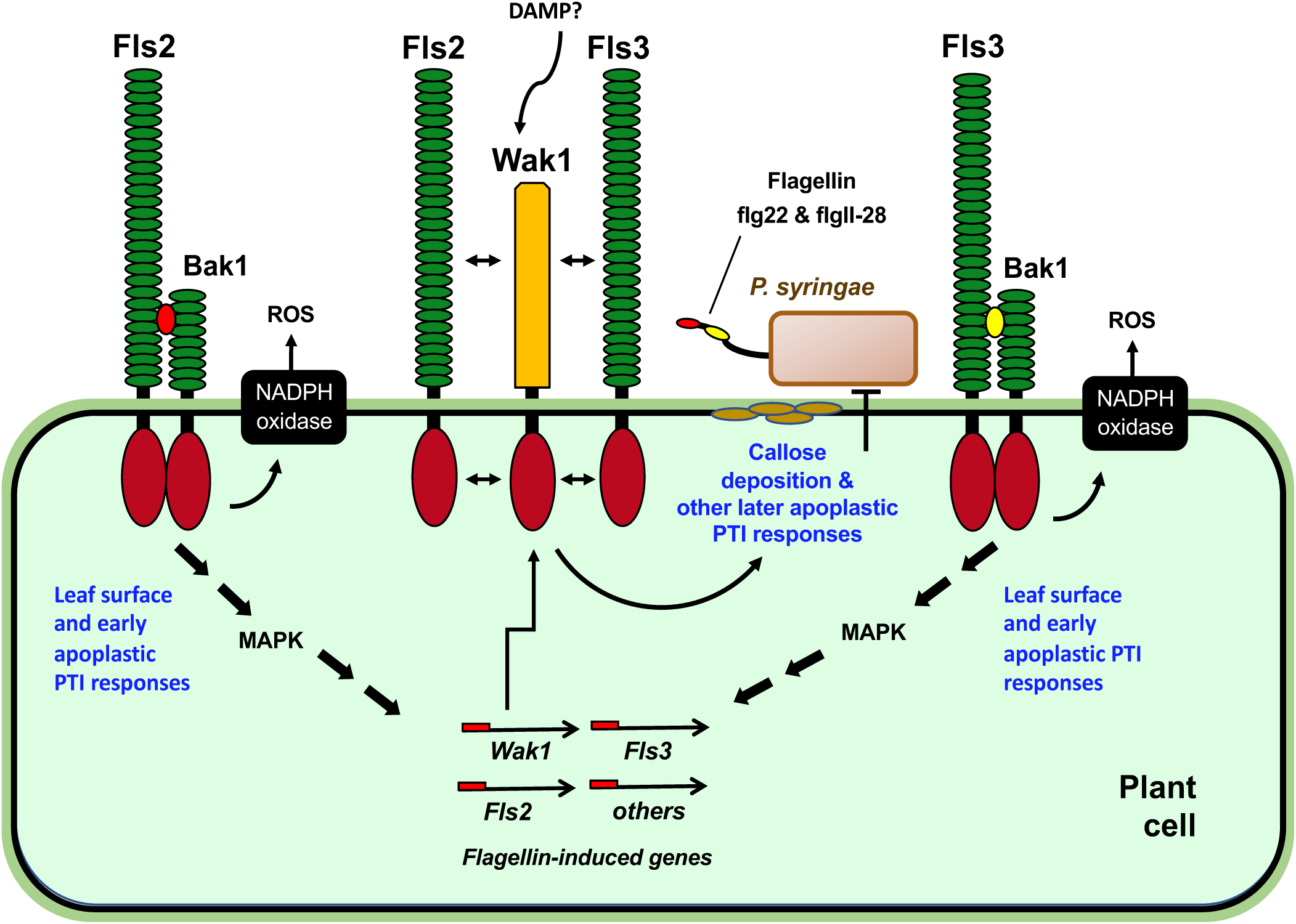
A model for SlWak1-mediated immunity in PTI. Transcript abundance of *Wak1* increases in mesophyll cells upon activation of the Fls2 or Fls3 pathways. This leads to increased accumulation of Wak1 protein which is then localized to the cell wall where it joins a complex containing Fls2 and Fls3. Wak1 might act as a scaffold and could be a receptor for a damage-associated molecular pattern (DAMP). Wak1 functions to promote deposition of callose at the cell wall and other immune responses that inhibit multiplication of the pathogen. See text for a full discussion of this model.

This model gives rise to several questions that will need to be addressed in the future. First, why is Wak1 not active in plant cells on the leaf surface, including stomata, but only functions when *Pst* enters the apoplastic space? This could be due to lack of *Wak1* gene expression, protein accumulation, association with the Fls2/Fls3 complex, or kinase activity in leaf surface cells. Second, how does Wak1 affect the cell wall-associated Fls2/Fls3 complex and is its activity in this complex influenced by perception of a DAMP? In Arabidopsis, AtWAK1 was demonstrated to bind pectin and OGs *in vitro* (Kohorn *et al*., 2009) and identified as the receptor for OGs *in vivo* (Brutus *et al*., 2010). Does Wak1 bind OGs and, if so, do OGs impact the way Wak1 associates with Fls2/3 and its role in PTI? Finally, it will be interesting to investigate possible differences in the transcriptome, metabolome, and proteome of the Δwak1 mutants in comparison with wild-type plants to understand what are the later PTI responses to which Wak1 contributes.

## Acknowledgments

We thank Robyn Roberts for comments on the manuscript and for sharing seeds of the Δfls2.1/fls2.2/fls3 line, Holly M. Roberts for assistance with vacuum infiltration and sequencing of the F2 populations, Brian Bell and Jay Miller for plant care, Joyce Van Eck for tomato transformation and Diana Lauff for assisting with experiments involving microscopy. Funding was provided by National Science Foundation grant IOS-1546625 (GBM) and Agencia Nacional de Promoción Científica y Técnica - Argentina PICT 2017-0916 (MAP).

## Author contributions

GBM and NZ conceived and designed the experiments. MP and HR performed the callose deposition assay and stomata measurements. NZ designed gRNAs, constructed vectors, performed genotyping and phenotyping experiments, and analyzed the data. NZ and GBM interpreted the data and wrote the manuscript. All the authors read and approved of the manuscript.

## Supplemental information

**Figure S1.**
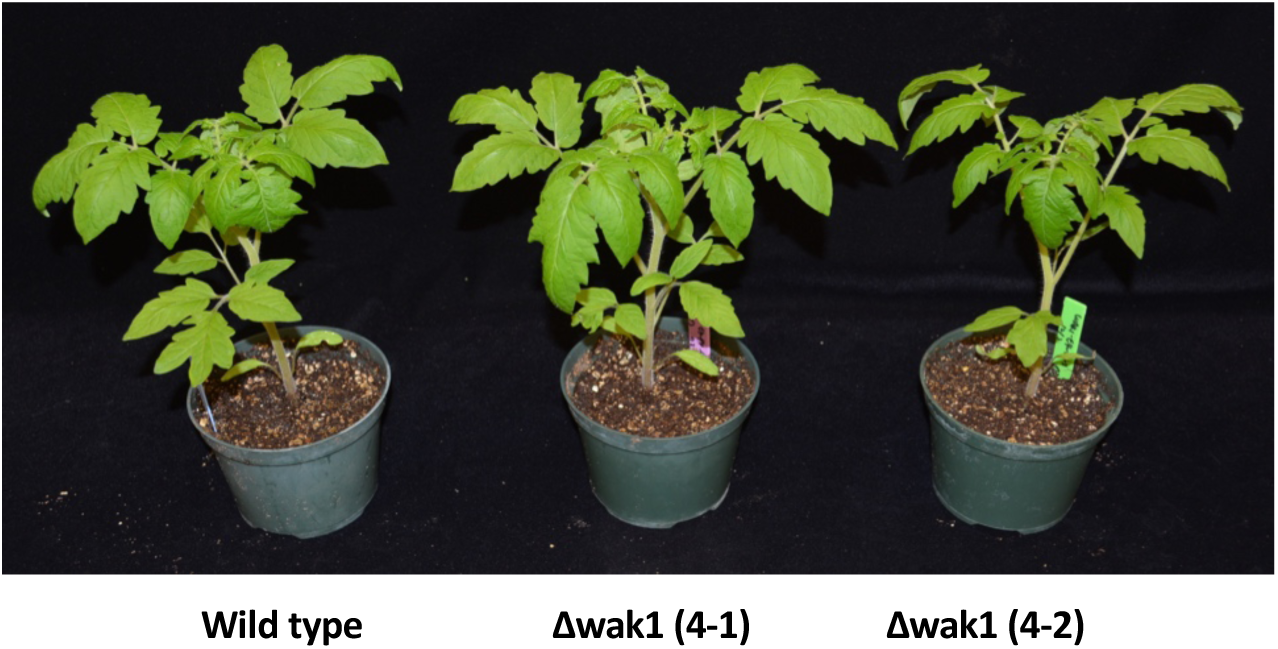
The growth, development and morphology of *Wak1* plants were indistinguishable from wild-type RG-PtoR plants. Photographs of four week-old wild-type RG-PtoR and the two Δwak1 mutant lines (4-1 and 4-2) grown in the greenhouse.

**Figure S2.**
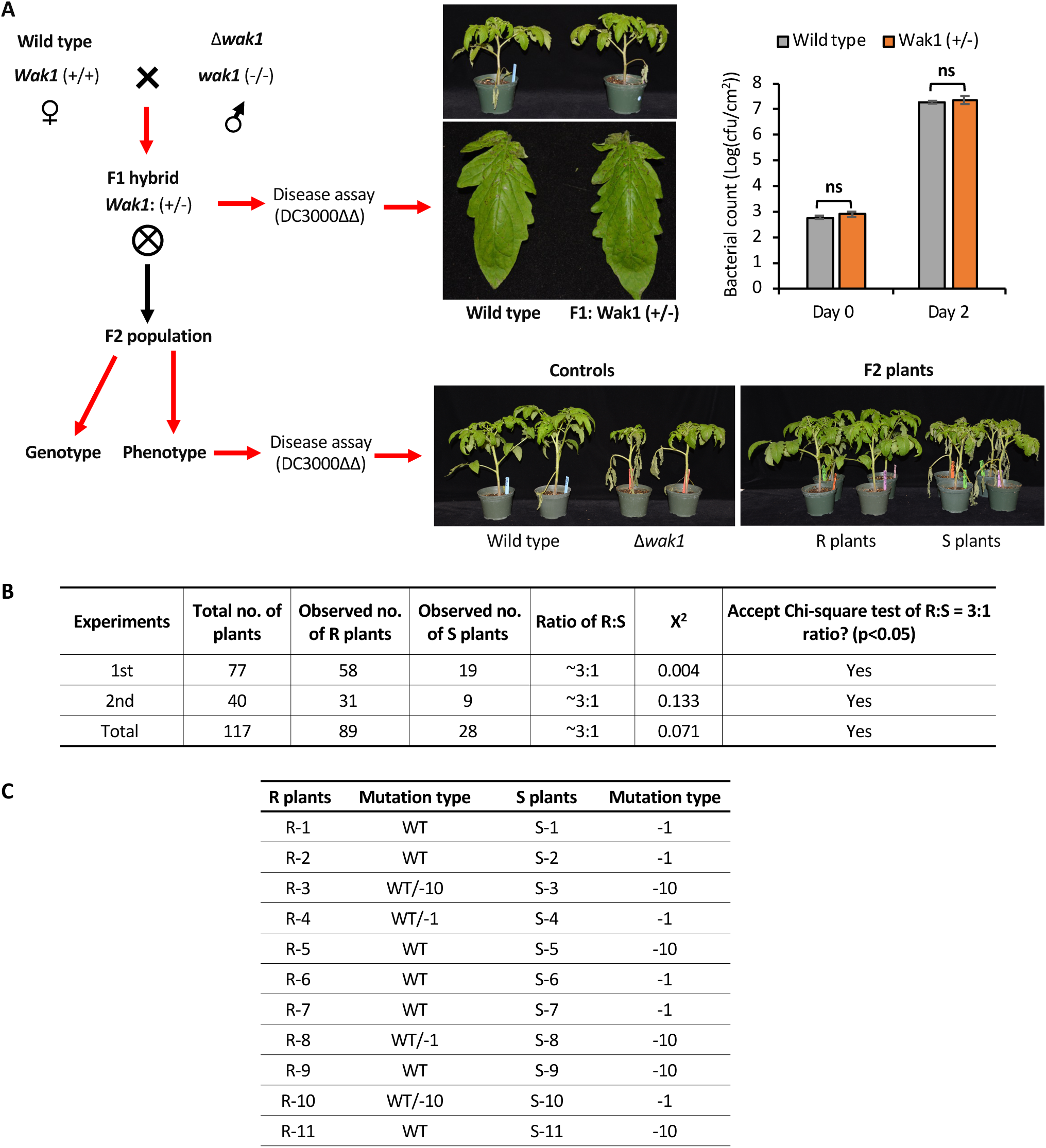
Enhanced susceptibility to DC3000ΔΔ co-segregates with the *wak1* mutations. **A**) The Δwak1 lines (4-1 and 4-2) were backcrossed to the wild-type RG-PtoR. Seeds from F1 hybrids were genotyped to confirm the heterozygous genotype (either WT/-1 bp or -WT/-10 bp; WT, wild type). F1 hybrids were tested with DC3000ΔΔ (as in Fig. 2A) and no significant difference was observed in disease symptoms or bacterial populations between F1 and wild-type plants. F1s were then selfed and F2 populations along with wild type (resistant control) and Δwak1 plants (susceptible control) were inoculated with DC3000ΔΔ. Number of resistant (R) and susceptible (S) plants in F2 populations were recorded. Photographs of disease symptoms were taken 5 dpi. **B**) Summary of disease assay with F2 populations. Chi-square test supported a segregation ratio of 3:1 (R:S) in the F2 population. **C**) Genotypes of representative resistant (11 plants) and susceptible plants (11 plants) by PCR and sequencing. F2 plants resistant to DC3000ΔΔ were either wild type or heterozygous mutants (either WT/-1 bp or WT/-10 bp), while all of the susceptible plants were homozygous mutants (either -1 bp or -10 bp). -#, # of base pair deletion in *Wak1.*WT, wild type.

**Figure S3.**
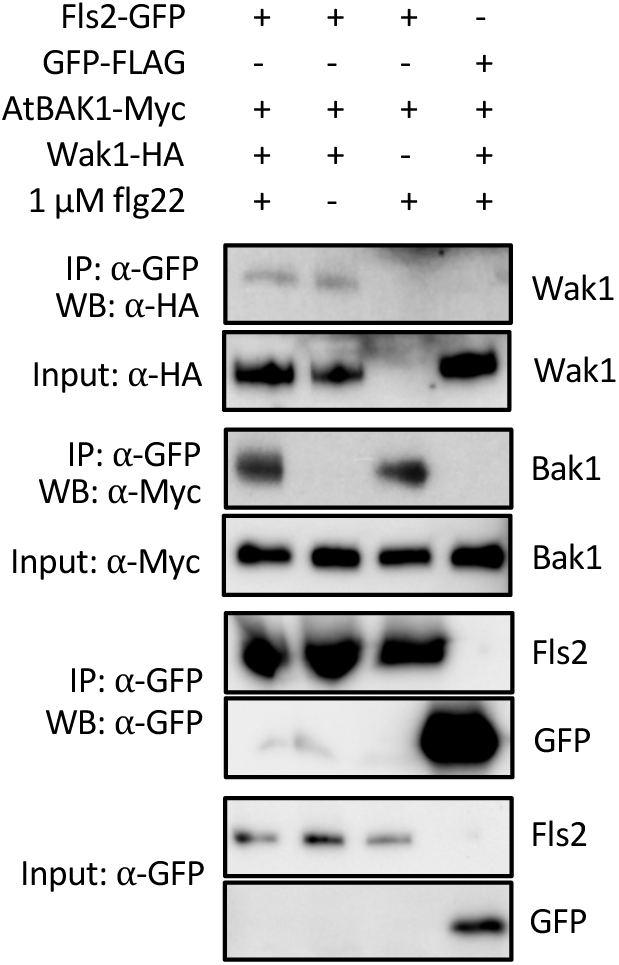
Wak1 associates with Fls2 independently of flg22. Proteins were extracted from *N. benthamiana* leaves expressing Fls2-GFP in combination with BAK1-Myc and/or Wak1-HA after treatment with or without 1 μM flg22 for 2 min and were used for immunoprecipitation using anti-GFP magnetic agarose beads. Wak1 was pulled down with Fls2 after treatment with or without flg22. Wak1, BAK1, Fls2, and GFP proteins were detected by immunoblotting with ⍺-HA, ⍺-Myc, or ⍺-GFP antibodies. This experiment was repeated twice with similar results.

**Table S1.**
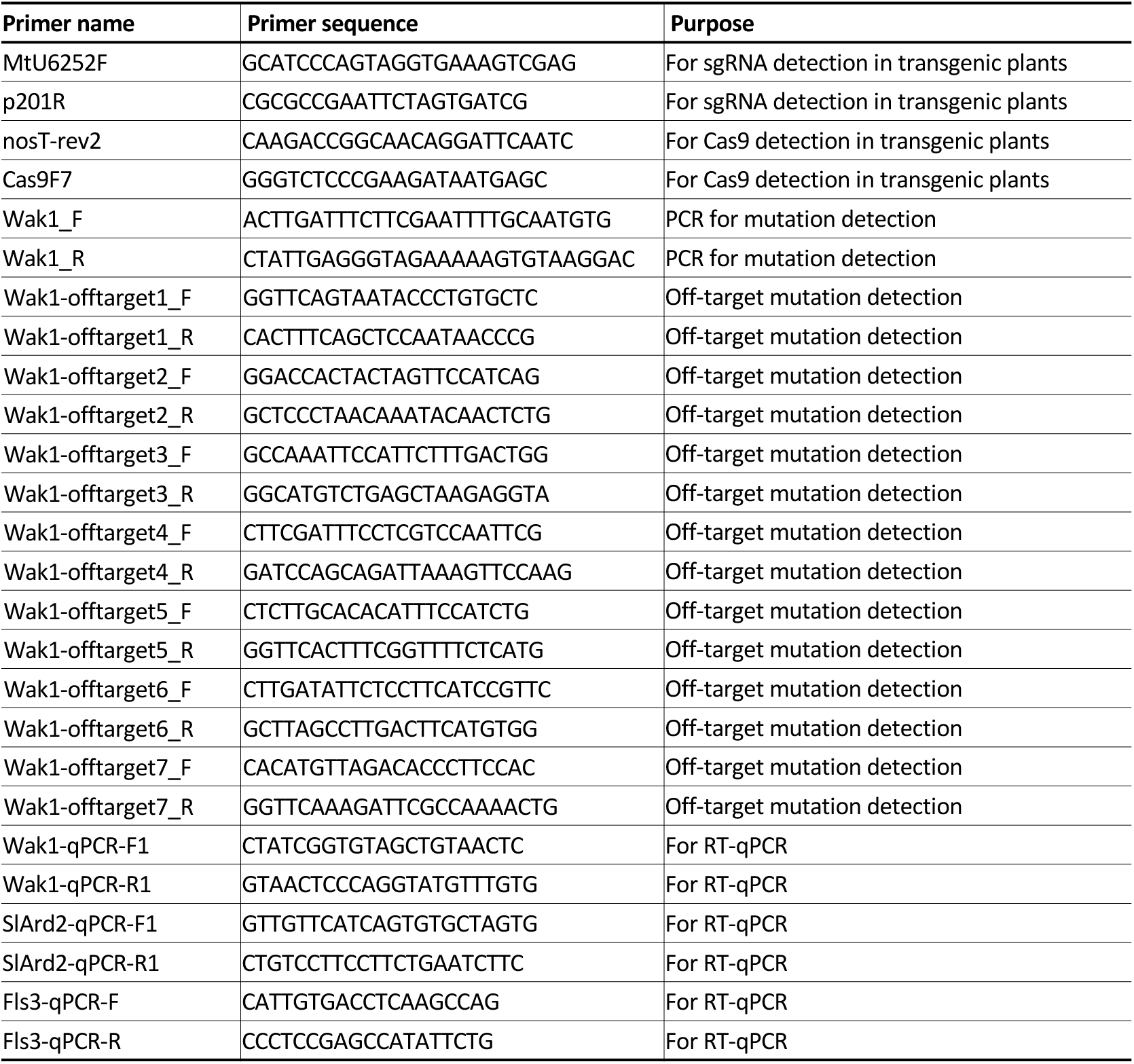
Primers used in this study.

